# Increasing threat: *Rhynchophorus ferrugineus* invaded mainland America through Uruguay

**DOI:** 10.64898/2026.09.11.750795

**Authors:** Martin Bollazzi, Vitor Pacheco da Silva, Sebastián Pita, Julián Sabattini, Andrea Listre, Carolina Negrone C, Federico Lopez Romanelli, Aymer A. Vásquez-Ordóñez

**Affiliations:** Entomología, Facultad de Agronomía, Universidad de la República, Montevideo, Uruguay; Entomología, Facultad de Ciencias, Universidad de la República, Montevideo, Uruguay; Genética Evolutiva, Facultad de Ciencias, Universidad de la República, Montevideo, Uruguay; Facultad de Ciencias Agrarias, Universidad Nacional de Entre Ríos CONICET, Paraná, Argentina; Laboratorio Biológico, Dirección General de Servicios Agrícolas, MGAP, Montevideo, Uruguay; Intendencia de Canelones, Canelones, Uruguay; Instituto de Ecología, Universidad Nacional Autónoma de México, México, México

**Keywords:** *Rhynchophorus ferrugineus*, invasion, America, Uruguay, weevil, pest, palms

## Abstract

*Rhynchophorus ferrugineus* (Olivier) (Coleoptera: Curculionidae) is one of the most dangerous palm pests worldwide. Here we report the invasion process of *R. ferrugineus* in Uruguay. Beyond morphological and molecular identification, we infer the geographic origin of the invasive population, describe the succession of introduced and native palms attacked, provide an easy-to-use key to differentiate the invader *R. ferrugineus* from the native *Rhynchophorus palmarum*, and model the ecological niche of *R. ferrugineus* in Uruguay and neighboring countries. First detected in 2022, the Uruguayan population of the weevil is the first to establish on mainland America, following the earlier island records from Curaçao and Aruba. Phylogenetic analysis showed that the Uruguayan *R. ferrugineus* sequences clustered within a single clade, most closely related to sequences from southern China, and clearly distinct from the sequences of Curaçao and Aruba, indicating at least two independent introduction events into the Americas from different source regions. After four years, *R. ferrugineus* has spread across an area of nearly 80,000 square kilometers and in January 2026 it reached Argentina, invading two points near the border with Uruguay. Over the first two years after the invasion, attacks focused on *Phoenix canariensis*, yet the three most common native palms in Uruguay, *Butia odorata*, *Butia yatay*, and *Syagrus romanzoffiana*, are currently being attacked. The diagnostic key to differentiate the pair *R. palmarum / R. ferrugineus* is based on three characters: the tip of the rostrum, the tegminal plate, and the ratio between the interocular space and the width of the rostrum at the base. Niche models showed that Uruguay and the neighboring areas of Argentina and Brazil are suitable for *R. ferrugineus*, and these are precisely the areas where both the highly susceptible *P. canariensis* and the native *B. odorata*, *B. yatay*, and *S. romanzoffiana* are present. Neighboring countries should therefore be prepared for the expansion of *R. ferrugineus* in the short term.

## Introduction

*Rhynchophorus ferrugineus* (Olivier) (Coleoptera: Curculionidae) is one of the most dangerous pests for palms worldwide because of its high ensuing mortality rate (1, 2). Its larvae feed on the meristematic tissue of the palm crown or of offshoots, causing damage that either kills the palm directly or facilitates the entry of lethal pathogens and secondary insect pests (1, 3, 4). *R. ferrugineus* has been reported attacking at least 40 species of palm trees (2), such as *Cocos nucifera* L., *Phoenix canariensis* Chabaud, and *P. dactylifera* L., in its native and invasive regions (1, 4–8). *P. canariensis* colonization represents a recent association (2), demonstrating its ability to colonize new hosts. For these reasons, it has been categorized as a quarantine pest in several countries in Africa, America, Asia, and Europe (9). In the last four decades, its invasive capacity has allowed it to colonize all continents except mainland America (2).

Molecular and historical evidence suggest that *R. ferrugineus* is native to Southeast Asia (1, 10). Travel, trade, and transportation are the main drivers of the rapid intercontinental geographic expansion of *R. ferrugineus* (11), with ornamental palm trade contributing significantly (1). In 1985, it was reported, for the first time, in the Middle East, very likely due to transport of palm trees from Asia (2, 4). Afterwards, this weevil has been reported in the Mediterranean, Canary Islands, North Africa, Japan, China, Taiwan, and the Caribbean Islands (1, 2). The phylogenetic reconstructions suggest possible geographic regions from which the invasive individuals originated, with Pakistan pointed as the origin of the populations that invaded the Middle East, and Thailand, Malaysia and southern China as the origin of those that invaded the Mediterranean (10). To date, *R. ferrugineus* has not been reported in mainland America. The closest records come from the Caribbean islands of Curaçao and Aruba. The Curaçao population probably originated from Egypt or the Mediterranean Basin, and the Aruban one from Curaçao, with the differences between them reflecting either post-introduction mutation or a second introduction event from Egypt, the Mediterranean Basin, or the native range in Northeast Asia (10).

Environmental niche analyses suggest that much of the American continent has favorable climatic conditions for *R. ferrugineus*, including South America (12, 13). The negative consequences of this weevil’s presence in mainland America could be significant, given its potential impact on native and commercial palms (14–16). In addition, such impacts could add to or synergize with the damage already generated by other palm tree borer-weevils, such as *R. palmarum* (*17*) and *Dynamis borassi* Fabricius (18).

Taxonomic confirmation of *R. ferrugineus* is paramount for rapid preventive management. However, some morphological variation occurs among species of this genus in diagnostic features such as coloration (10, 19, 20), endophallus armature (2, 21), concavity of subgenal sutures (19, 22), and pronotal shape (10, 22). In America, misidentification between the native *Rhynchophorus palmarum* (L.) and the invasive *R. ferrugineus* should be particularly avoided, given the similar attack symptoms, confusing reddish coloration of *R. palmarum* (19), and documented variability in some classical morphological characters for taxonomic differentiation shown by *R. palmarum*, such as shape of the pronotum, mandible, and subgular suture (3, 23). Molecular analysis with the mitochondrial gene cytochrome c oxidase I (COI) has allowed a very accurate differentiation of *R. ferrugineus* from five of the eight species currently recognized in the genus, namely *R. vulneratus* (Panzer), *R. bilineatus* (Montrouzier), *R. cruentatus* (Fabricius), *R. phoenicis* (Fabricius), and *R. palmarum* (L.) (5, 10, 19). For example, this made it possible to determine that the species that invaded California in the United States corresponded to *R. vulneratus* and not, as initially reported, to *R. ferrugineus* (10).

This work is aimed at investigating the invasion process of *R. ferrugineus* in Uruguay, the first country in mainland America reached by this weevil. Beyond morphological and molecular identification, we infer the geographic origin of the invasive population, describe the succession of introduced and native palms attacked in recent years, and develop an easy-to-use diagnostic morphological key for the pair *R. ferrugineus/R. palmarum*. Finally, we model the ecological niche of *R. ferrugineus* to generate a regional suitability map and identify areas where the weevil could expand from Uruguay by spreading across susceptible native palms.

## Methods

### Morphological identification

Between March and July 2022, thirty-one specimens, sixteen females and fifteen males presumably belonging to *R. ferrugineus,* were analyzed. All specimens were collected manually from the crown of *P. canariensis* palm trees. Material examined: Uruguay: Canelones, Santa Lucia, 2022, 3 ♀ and 1 ♂ (34°27′01.74″S 56°22′52.23″W); Canelones, Estación Margat, 2022, 2 ♂ (34°28′52.71″S 56°20′40.15″W); Canelones, Prado, 2022, 8 ♀ and 4 ♂ (34°31′37.14″S 56°16′26.67″W). San José, Colonia Wilson, 2022, 3 ♀ (34°42′10.20″S 56°31′25.43″W). Florida, Mendoza, 2022, 2 ♂ (34°17′00.68″S 56°12′49.35″W). Montevideo, Lezica, 2022, 2 ♀ and 6 ♂ (34°47′46.30″S 56°14′57.02″W).

Taxonomic identification was based on direct observation using an Olympus SZX16 and a Nikon SMZ1270 stereomicroscope. Eight dissections of male genitalia were performed to confirm the diagnosis of the species. The genitalia were cleaned in KOH 10%, rinsed in distilled water, and subsequently mounted in glycerin to identify the morphological diagnostic characters. Morphological characterization of *R. ferrugineus* was based on previous taxonomic studies (3, 22–24).

In addition to species determination, we applied morphological characterization to develop an easy-to-use diagnostic morphological key for the pair *R. ferrugineus/R. palmarum.* For that, we expanded the characterization of the morphological variation made by Löhr et al. (19), by working with 65 more exemplars of *R. palmarum* collected in Argentina, Brazil, Colombia, Guatemala, México and Nicaragua (Museo de Zoología, Facultad de Ciencias, and Colección Nacional de Insectos of the Instituto de Biología, Universidad Nacional Autónoma de México, México City) (Supplementary Table 1). As a result, morphological variation in *R. palmarum* characters important for distinguishing the *R. ferrugineus/R. palmarum* pair was assessed in 520 *R. palmarum* exemplars to date: 65 during this study added to the 455 from Löhr et al. (19). The diagnostic characters of those exemplars were contrasted to the 31 *R. ferrugineus* collected in Uruguay in 2022 to determine the invader species, following the classical characters proposed by Giblin Davis et al. (3) and Wattanapongsiri (23), yet considering the morphological variation of *R. palmarum* described in Löhr et al. (19).

### Molecular analysis

Molecular analyses of the mitochondrial *cytochrome c oxidase I* (*COI*) gene were conducted with two aims: to confirm the morphological identification of the specimens collected in Uruguay, and to infer the geographic origin of the invasive population, including whether it shares a haplotype with the only previous *R. ferrugineus* record for the Americas, in Aruba and Curaçao (10).

DNA was extracted from the legs of 13 adult *R. ferrugineus* beetles collected in Uruguay, following the protocol by Medrano et al. (25) with slight modifications, which uses NaCl 5M for protein precipitation. PCR was carried out to amplify a fragment of the *COI* gene using the conserved primers LCO1490 and HCO2198 from Folmer et al. (26). PCR products were evaluated using agarose gel electrophoresis. Amplified *COI* PCR products were sent to Macrogen Inc. (Seoul, Korea and sequencing on both strands. The generated files in ab1 format were used as input in the R package *sangeranalyseR* (27) to perform trimming by quality and the generation of a consensus sequence for each individual. A BLAST search was conducted to compare the consensus sequences with the NCBI standard nr database.

A phylogenetic tree was estimated by maximum likelihood (ML) with the R package *phangorn* (28). The tree was bootstrapped 100 times to assess node support. To find the best-fitting substitution model, the *phangorn* JModeltest function was used. Model selectionion was based on the Bayesian Information Criterion (BIC). Tree visualization and editing were performed using the R package *ggtree* (29).

### Ecological modeling and potential expansion from Uruguay

We modeled the *R. ferrugineus* ecological niche using global occurrence data to assess the distribution potential for the red palm weevil in Uruguay and neighboring countries. We compared the suitability of *R. ferrugineus* occurrence with native palm distributions shared with Argentina and Brazil.

Presence records of the red palm weevil were collected from the Global Biodiversity Information Facility (http://www.gbif.org/species) database, and complemented with a review of the scientific literature (12) and cross-validated against the specialized invasive-pest databases EPPO (https://gd.eppo.int/) and CABI (https://www.cabidigitallibrary.org/). To avoid pseudoreplication of local environments owing to artificial clustering of occurrence sites, we reduced the dataset such that each observation fell inside a separate 20 km grid cell, leading to a total of 429 distinct occurrence records for the red palm weevil, including the new records from Uruguay and Argentina (Supplementary Table 2) (30, 31). We downloaded the 19 bioclimatic variables from the WorldClim database version 2 (http://www.worldclim.org/), averaged for the 1970–2000 period, at a spatial resolution of 2.5 minutes, approximately 20-kilometer resolution at the equator (32). Of these, we chose the seven that are relatively uncorrelated globally (33): annual mean temperature, mean diurnal temperature range, maximum temperature of the warmest quarter, minimum temperature of the coldest quarter, annual precipitation, precipitation of the wettest quarter, and precipitation of the driest quarter (34). The selected geographical distribution records and bioclimatic variables were imported into the MaxEnt model to analyze and generate a regional and worldwide suitability map for *R. ferrugineus* distribution.

## Results

### First reports and current distribution in Uruguay

*Rhynchophorus ferrugineus* was first detected in a public park in Canelones and in the city of Santa Lucía, two sites approximately 13 km apart, between February and April 2022 (Figure 1**A**). Detection was visual, prompted by the symptoms shown by ornamental *P. canariensis* palms (Figure 1**B**, **C**). Adults and larvae were collected from the crowns after they had fallen, once the leaves started detaching. The affected area was located 40-50 km north of Montevideo, the main port of entry to Uruguay. During 2022 and 2023, *R. ferrugineus* became well established in Montevideo, the capital city. Attacks focused primarily on *P. canariensis*. Since then, other introduced ornamental palms, including *Washingtonia robusta* H.Wendl, *Washingtonia filifera* (Gloner ex Kerch., Burv., Pynaert, Rodigas & Hull) de Bary*, and Trachycarpus fortunei* (Hook.) H.Wendl., also began to be attacked.

**Figure 1:**
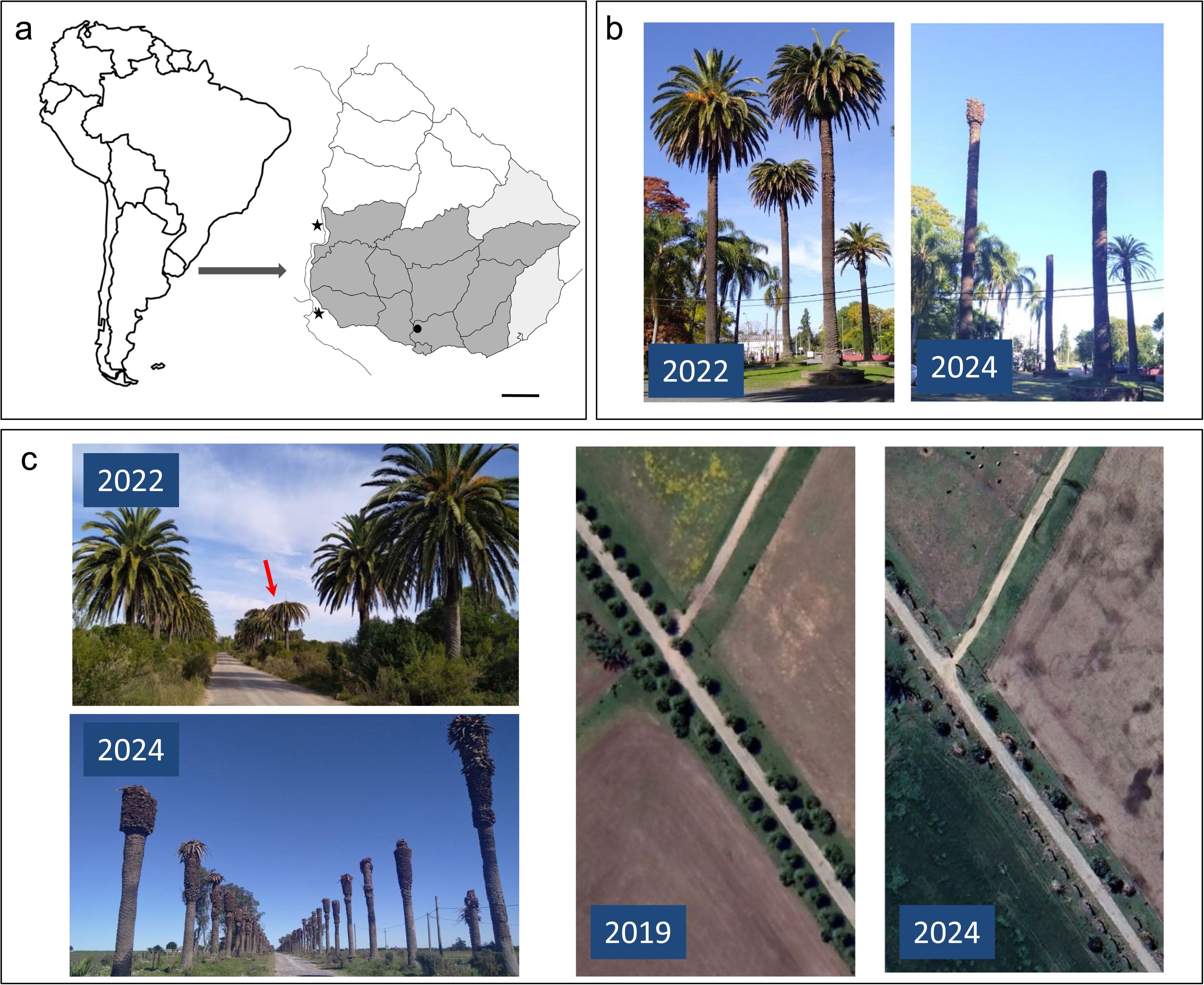
Invasion process of Uruguay by *Rhynchophorus ferrugineus* since 2022. (**A**) South America and Uruguay map indicating the location of the first report of *Rhynchophorus ferrugineus* (black dot) in 2022. Stars represent the two recent invasions in Argentina. For Uruguay, its current distribution is shown by the grey-filled states. Dark grey: palms have been confirmed to be killed by *R. ferrugineus*. Light grey: only the weevil was captured, yet dead palms have not yet been confirmed. Scale bar: 50 km (**B**) *Phoenix canariensis* in Prado Park, Canelones, Uruguay. In 2022, a group of three *P. canariensis,* more than a hundred years old, started being attacked. While two palms showed the typical symptoms by May 2022, the third was still without evident signs of attack. In April 2024, all palms were killed by *R. ferrugineus* and dozens of palms in the surroundings. At the back, *Syagrus romanzoffiana* and *Butia odorata* palms remain unattacked. (**C**) At the beginning of the XX century*, P. canariensis* was planted as an ornamental palm on several portions of the roads in Uruguay. In this case, a portion of more than 4km long with hundreds of palms, Paso Pache, Florida, Uruguay. In March 2022, the first *P. canariensis* showed the typical generalized crown decay caused by *R. ferrugineus* (red arrow). In 2024, all palms were dead. Satellite pictures on the right show comparatively the same portion of the road in 2019, three years before the arrival of *R. ferrugineus*, and in 2024, when few palms were still alive. Pictures by M. Bollazzi. Google Earth Pro 7.3.6.10201 (64-bit) (34°20′25.16″S 56°14′48.18″W).

Between the second half of 2023 and early 2024, several reports indicated that *R. ferrugineus* had begun to attack native hosts (Figure 4**A**). In April and May 2024, *R. ferrugineus* was collected from dead *B. odorata* Noblick palms in Montevideo. Simultaneously, larvae were collected from the native *S. romanzoffiana* Glassman palm in Canelones city, near the location of the first reports from 2022. Molecular methods later confirmed these larvae as *R. ferrugineus*. Mortality of *B. yatay* Becc. palms, the third native palm attacked by *R. ferrugineus*, was later confirmed.

Four years after the confirmed start of the invasion, *R. ferrugineus* has spread across nearly 80000 km², approximately 370 km from west to east and 240 km north from the southern coast (Figure 1**A**). At present, the true expansion of *R. ferrugineus* in Uruguay is probably underestimated. In January 2026, *R. ferrugineus* arrived in Argentina, where it is still restricted to the regions bordering Uruguay (Figure 1**A**). It was first recorded attacking a *P. canariensis* palm on Martín García Island (30), in the Río de la Plata, the natural frontier between Argentina and Uruguay. The island lies 4 km from the nearest point on the Uruguayan coast. In August 2026, *R. ferrugineus* was also detected in the Entre Ríos province (31), 170 km north of Martín García Island. This probably represents a second introduction event from Uruguay, since *R. ferrugineus* individuals were collected inside an exemplar of the native palm *S. romanzoffiana* located near the Uruguay River coast. *R. ferrugineus* is already present in Uruguay less than 10 km away, on the opposite bank of the river.

### Morphology

Morphological characterization confirmed that the collected exemplars in Uruguay since 2022 belong to *R. ferrugineus*. The morphological comparison of the 65 *R. palmarum* specimens of South and Central America performed in this work, together with the *R. ferrugineus* from Uruguay, confirmed the suitability of the three candidate diagnostic characters to differentiate the two species: two proposed by Löhr et al. (19) and others (3, 22–24), and one confirmed here. These characters are: the tip of the rostrum, the ratio between the interocular space and the width of the rostrum at the base, and tegminal plate (Figure 2). We did not find a clear difference in pronotum shape as an easy-to-use diagnostic character to differentiate *R. ferrugineus* from *R. palmarum* (3, 21), consistent with the results of Hallet (22) and Rugman Jones (10). We therefore did not include this character in our diagnostic key until a more extensive revision of the *Rhynchophorus* genus is performed. Nonetheless, differences in the male tegminal plate shape were consistent, as suggested by Wattanapongsiri (23) and Löhr et al. (19), and we propose to consider it as a diagnostic character.

**Figure 2:**
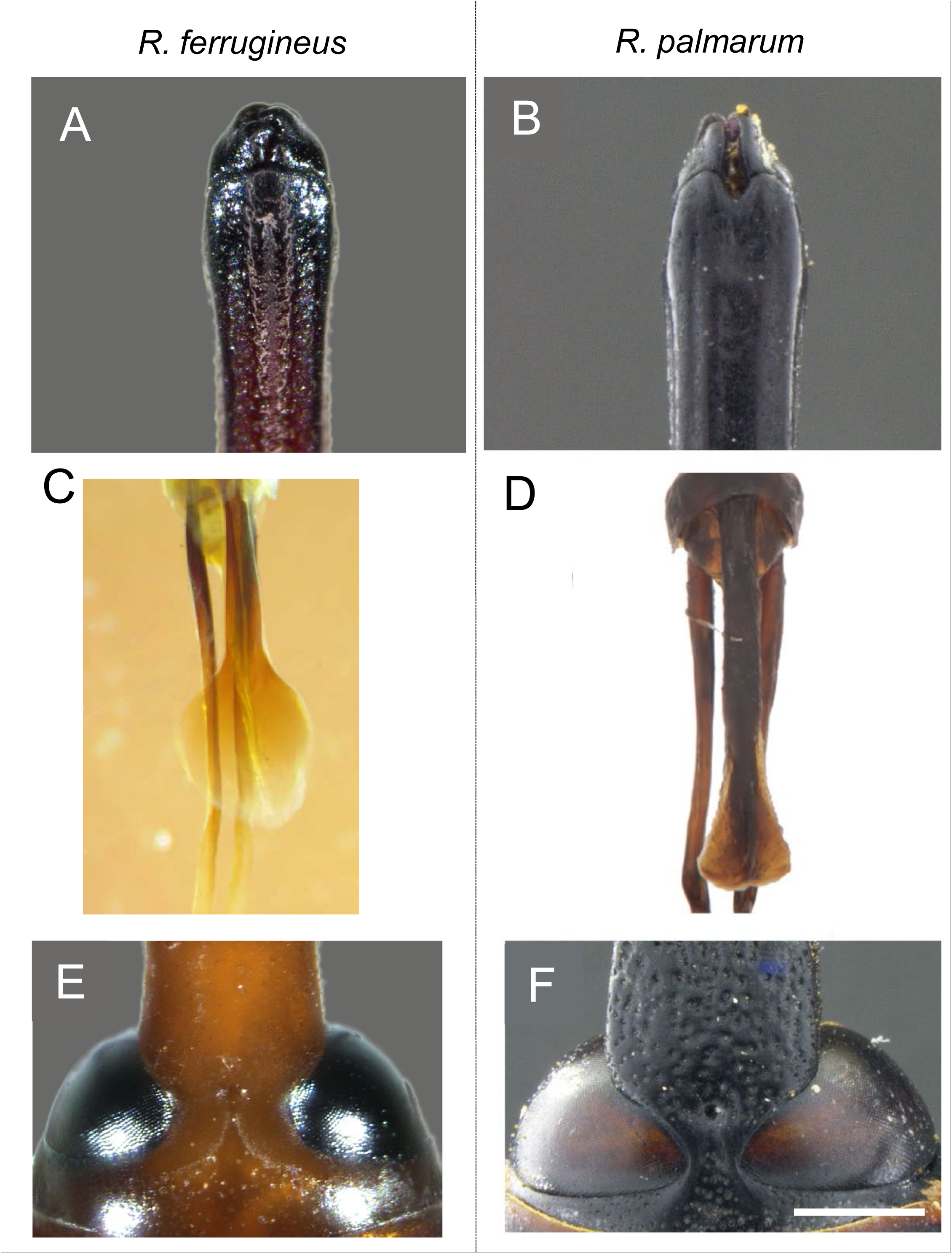
Characters for taxonomic differentiation between the invader *Rhynchophorus ferrugineus* and the native weevil *R. palmarum.* Tip of rostrum dorsally (**A**, **B**), tegminal plate (**C**, **D**), and interocular space and rostrum at the base (**E**, **F**) (dotted lines). Scale bar 1 mm. Pictures by M. Bollazzi, A. Listre y A. Vazquez-Ordoñez.

### Key to the South and Central American species of *Rhynchophorus* of economic importance (adults)

1a. Tip of rostrum not grooved but oval distally (Figure 2**A**); male genitalia: tegminal plate is round and flat, being from circular-shaped to fan-shaped plate (Figure 2**C**); interocular space between 0.54 and 0.64 times the width of the rostrum at the base (Figure 2**E**); *R. ferrugineus* 1b. Tip of rostrum dorsally grooved or nearly truncated (Figure 2**B**); male genitalia: tegminal plate is triangular (Figure 2**D**); interocular space between 0.12 and 0.35 times the width of the rostrum at the base (**Figure 2F**) *R. palmarum*

### Molecular marker determination

We obtained 13 barcode sequences of *R. ferrugineus* samples from Uruguay. Several sequences were retrieved from the BLAST search using Uruguayan sequences as query. Some hits were not included in the analyses due to their short length. A 375 base pair alignment was employed, including 258 sequences assigned to *R. ferrugineus.* Four haplotypes were observed from the 13 *R. ferrugineus* samples, with three segregating sites between them. In addition, sequences from *R. cruentatus* (2), *R. bilineatus* (2), and *R. vulneratus* (3) were added (Acc. Numbers in Supplementary Table 3). The ML tree is shown in Figure 3. The Uruguayan sequences clustered within a single clade, most closely related to *R. ferrugineus* sequences from southern China (35). This Uruguayan clade is clearly distinct from the Caribbean sequences of Aruba and Curaçao: Uruguayan sequences differ from Curaçao and Aruba by 10–11 and 11–12 substitutions out of 375 bp, respectively, compared to a single substitution separating Aruba from Curaçao.

**Figure 3:**
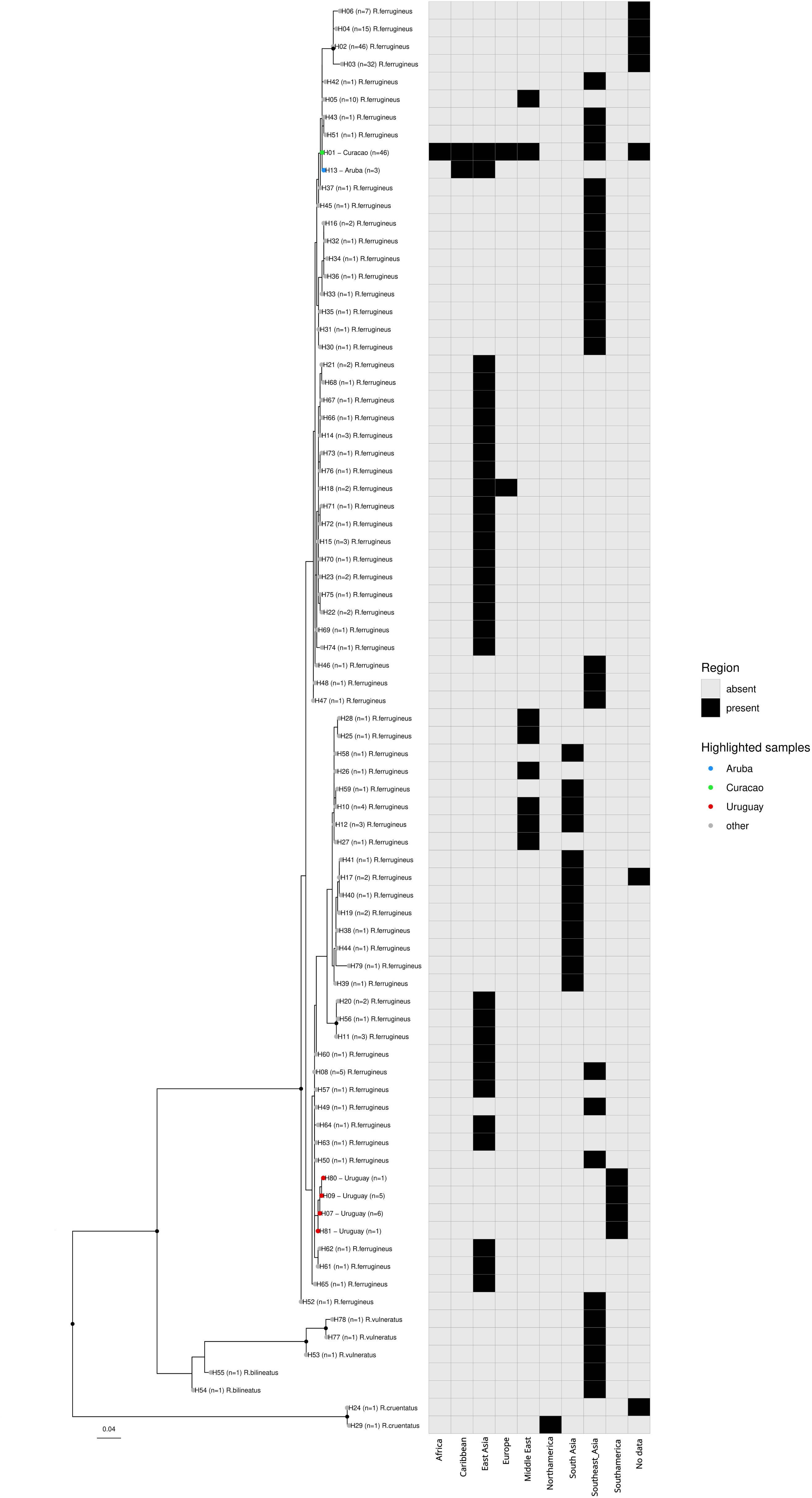
Maximum likelihood phylogenetic tree based on COI gene fragments for species of the genus *Rhynchophorus*. Each tip represents one haplotype; the number of specimens sharing it is given in parentheses (n). Colored tips highlight the haplotypes from Uruguay (this study, red), and from the only previous *R. ferrugineus* records for insular Americas, Aruba (blue) and Curaçao (green); gray tips correspond to all other GenBank haplotypes. Black dots on internal nodes indicate bootstrap support over 0.9. The grid on the right shows, for each haplotype, in which biogeographic region it has been recorded (black = present, gray = absent); “No data” indicates GenBank records with no country of origin reported. Scale bar: substitutions per site.

### Ecological modelling

The ecological niche model analysis showed a high area under the receiver operating characteristic curve (AUC > 0.9, training AUC = 0.970, test AUC = 0.969), indicating strong predictive performance (Supplementary Figure 1). The predicted habitat suitability for *R. ferrugineus* in Uruguay and the neighboring regions of Brazil and Argentina is shown in Figure 4**B**. The worldwide high-resolution map is given in Supplementary Figure 2. Near the southern Uruguayan coast, where the first attacks were reported in 2022, the model shows medium suitability. On the Argentinian side, the recently invaded region also shows intermediate predicted suitability for *R. ferrugineus*. *P. canariensis* palms, the most attacked host during the invasion of Uruguay, are commonly present in the areas of medium predicted suitability for *R. ferrugineus*. Moreover, the suitable areas overlap with the known distribution of the native palms currently being attacked by *R. ferrugineus.* The Maxent model shows how climatic factors influence the potential distribution of *R. ferrugineus*. Among all modeled bioclimatic variables, mean temperature from the coldest quarter, annual mean temperature, and mean diurnal range contributed most to the model (45.6, 26.0, and 20.0 %, respectively). All remaining variables explain less than 10% of the potential distribution (Supplementary Figure 1).

**Figure 4:**
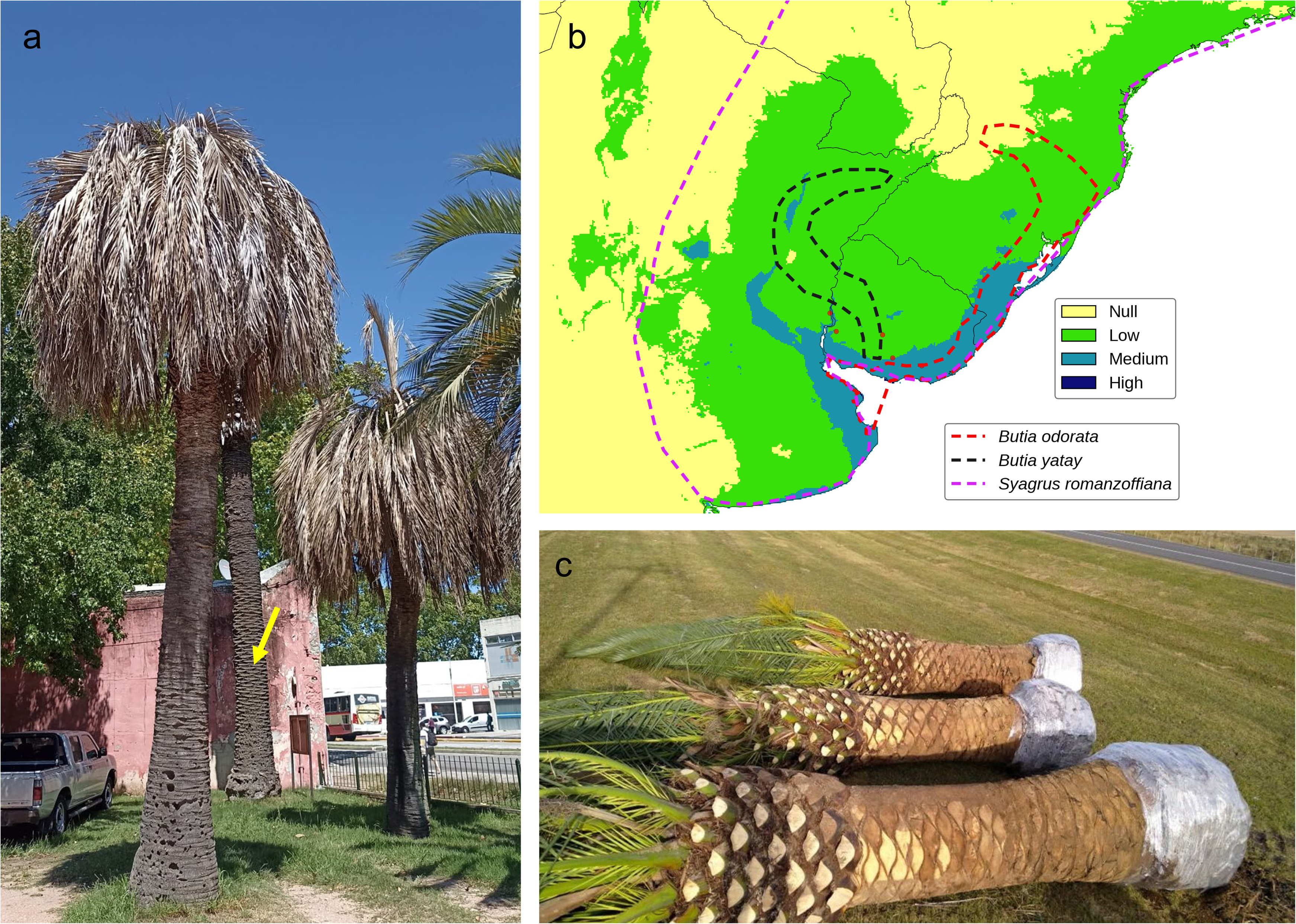
Niche analysis showing the potential distribution for *R. ferrugineus* in Southern South America compared to the distribution areas of the native palm hosts that *R. ferrugineus* has already attacked. (**A**) The first reports of native *Butia odorata* palms attacked by *Rhynchophorus ferrugineus* date from late 2023 and early 2024. Left side: two *B. odorata* palms with attack symptoms in a spot where a *Phoenix canariensis* was killed by *R. ferrugineus* a year before (the remaining *P. canariensis* trunk is marked with a yellow arrow at the back). (**B**) *R. ferrugineus* ecological niche modeling for Uruguay and portions of Argentina, Brazil, and Paraguay, superimposed on a schematic known distribution of the native palms *B. odorata*, *B. yatay,* and *Syagrus romanzofiana* (43, 55–57). (**C**) Three *P. canariensis palms* ready to be traded as ornamental palms beside a road in 2022. The picture was taken in the area where *R. ferrugineus* started attacking palms (black dot in Figure 1A). Pictures by M. Bollazzi.

## Discussion

The geographical distribution of *R. ferrugineus* has expanded to mainalnd America. This record adds to the report on Curaçao and Aruba islands for the Western Hemisphere (10). To test whether the Uruguayan population shares a haplotype with this earlier Caribbean invasion, we added the COI sequences from Aruba and Curaçao deposited by Rugman-Jones et al. (10) (GenBank KF311433 and KF311454-458, respectively) to our analysis. Consistent with Rugman-Jones et al. (10), the five Curaçao sequences were identical to each other and differed from the single Aruba sequence by a single nucleotide substitution. In contrast, all 13 *R. ferrugineus* sequences from Uruguay differed from both Curaçao and Aruba by 10 to 12 substitutions out of 375 bp, and are clustered with sequences from southern China (35). This indicates that the Uruguayan and Caribbean populations of *R. ferrugineus* do not share a haplotype and most likely represent at least two independent introduction events into the Americas, from different source regions within the native range: most likely Thailand/Malaysia for the Caribbean (10), and China for Uruguay. Asia is the region that traded the most with Uruguay (36), which could have facilitated this second introduction event. Whether *R. ferrugineus* entered Uruguay inside a palm host remains an open question, and the introduction of adults cannot be ruled out either, since ports are a focus of invasive coleopterans worldwide (37, 38).

At the time of the first reports in 2022, *R. ferrugineus* was detected almost simultaneously at two locations 55 km apart near Montevideo city. This rapid initial geographical expansion could be explained by the weevil’s capacity to fly up to 60 km in 24 hours (39). Moreover, this weevil likely arrived before 2022, as many attacked palms were already reported at both locations, suggesting the presence of more than one generation. The life cycle of this species takes 45 to 180 days (40). Given the identical symptoms shown by *P. canariensis* when attacked by both the exotic *R. ferrugineus* and the native *R. palmarum*, early attacks by *R. ferrugineus* were likely mistaken for those caused by *R. palmarum*. This likely hindered assessment of the red palm weevil’s spread, both in terms of the hosts attacked and its geographical extent. Several reports indicated an increase in dead *P. canariensis* palms from 2020–2021, before the presence of *R. ferrugineus* was confirmed. Thus, the exact time, place, and circumstances under which *R. ferrugineus* invaded mainland South America through Uruguay, which is acting as a bridge for the weevil to invade Argentina, remain unknown.

Invasion seems to have followed a two steps process since 2022. During the first two years, *R. ferrugineus* attacked mostly *P. canariensis* and other ornamental palms, and only from 2024 onwards the attacks focused on the native *B. odorata*, *B. yatay* and *S. romanzoffiana*. This pattern is similar to that reported in other areas (5, 41), with frequent initial attacks on the ornamental palm *P. canariensis* and less frequent attacks on other palm species. Of great concern, the three most common native palms in Uruguay and in the neighboring areas of Argentina and Brazil, *B. odorata, B. yatay*, and *S. romanzoffiana,* are being attacked. All three species have been reported as hosts of *R. ferrugineus* outside South America, where they are cultivated as ornamentals (1, 42). This association emphasizes the potential impact of this weevil on American native palms (Figure 4**A**, **B**), as *B. odorata and B. yatay* are key components of the *Butia* palm ecosystem, which is already of serious conservation concern (43). *R. ferrugineus* has been previously reported to complete its life cycle on *Butia* palms under laboratory conditions (44).This weevil is also likely to affect other species of the genus, such as *B. noblickii*, which is categorized as endangered according to the IUCN (45).

Currently, the main concern is its rapid geographic expansion toward neighboring Argentina and Brazil, similar to what has been reported in previous events in other countries (41, 46). This is supported by the niche analysis carried out in this study and those performed by Ge et al. (13) and Wang et al. (47). However, this contradicts the findings of Fiaboe et al. (12), likely because the current model used presence data from Uruguay. The risk of this pest invading other countries close to Uruguay is high. This risk arises not only from the suitability for *R. ferrugineus* shown by the niche models, but also from the predominance of the highly susceptible *P. canariensis* across Uruguay and its distribution associated with roads and cities leading into Brazil and Argentina. In addition, there is an overlap of the area already invaded by *R. ferrugineus*, mostly attacking *P. canariensis*, the natural corridors of suitability to Argentina and Brazil, and the distribution of the native hosts attacked by *R. ferrugineus*, such as *B. odorata*, *B. yatay,* and *S. romanzoffiana* (Figure 4**B**). Besides natural expansion, palm trade should also be considered. Uruguay was an exporter of *P. canariensis* as ornamental palms (Figure 4**C**), although its transport and commercialization are now forbidden. Nevertheless, smuggling of *P. canariensis* adult palms, each worth several thousand US dollars, across land borders without natural barriers, to Brazil for instance, makes exhaustive controls extremely difficult. Brazilian researchers have already reported the introduction of *R. ferrugineus* to Brazil from Uruguay (48). The weevil was found infesting a *P. canariensis* in São Paulo state, in January 2022, on a palm that had been brought from Uruguay by land transport in December 2021. This single infected palm was not enough to establish an invader population, and no further sightings have been reported. This shows that a single infested palm can establish a focus more than 1000 kilometers away from the source area. It also has implications for the timing of the Uruguayan invasion. Palms leaving Uruguay were already infested by late 2021, months before the weevil was officially detected. *R. ferrugineus* was most likely present in Uruguay one to two years before its confirmation in 2022, which is consistent with the increase in dead *P. canariensis* reported since 2020–2021.

Management strategies aimed at mitigating the spread of *R. ferrugineus* in Uruguay are mainly focused on chemical control (49). Aside from the known pros and cons of insecticides for controlling the red palm weevil (50, 51), conditions leading to efficient management are currently lacking in Uruguay, such as knowledge of the length of the life cycle and the timing of flight peaks. Knowing the annual variation in life cycle stages is crucial for proper chemical control, since the efficacy of authorized insecticides and application methods depends on *R. ferrugineus* life stages (52, 53). Another factor is that although local governments reacted rapidly at the beginning of the invasion by implementing controls, the rapid spread and the fact that attacks occurred mainly on private properties drastically reduced the capabilities of those institutions to contain *R. ferrugineus.* This hinders regional management, since the decision to control lies in private hands, with costs for treatment and extraction that are almost impossible to afford for average families. This results in a rapid spread of *R. ferrugineus* that will not stop until the population of susceptible palms drastically decreases.

Neighboring countries should be prepared for the expansion of *R. ferrugineus* in the short term. Besides chemical control, which is a fundamental step in this process, an accurate taxonomic diagnosis is essential. This could be challenging for *R. ferrugineus* since, besides coloration, several diagnostic morphological characters are shared with *R. palmarum* (19), which is abundant in Uruguay, Argentina, and Brazil (17). Molecular analysis of the COI gene allows for accurate diagnosis (19); however, it can be a costly and time-consuming strategy. To provide authorities, researchers, and laypersons with an easy-to-use tool for morphological differentiation, we provide an updated key for differentiating the invasive *R. ferrugineus* from the native *R. palmarum*. To devise adequate containment measures, and to understand the cause behind the current crisis, it is necessary to evaluate the combined effects of agricultural management, host susceptibility, and climate variables on infestation levels of palms caused by *R. ferrugineus*, as has already been done for other South American pest weevils such as *R. palmarum* and *Dynamis borassi* (54).

## Conflict of Interest

The authors declare that the research was conducted in the absence of any commercial or financial relationships that could be construed as a potential conflict of interest.

## Author Contributions

Conceptualization: MB, VP, SP, JS, AV. Writing – original draft: MB, VP, SP, JS, AV. Writing – review & editing: MB, VP, SP, JS, AV. Data curation: AL, AV, SP, VP. Formal Analysis: MB, SP, JS, AV. Funding acquisition: MB, JS, AV, SP, VP. Investigation: MB, VP, SP, JS, AV, CN, FL. Methodology: MB, VP, SP, JS, AV, CN, FL. Project administration: MB. Resources: MB, CN, FL. Supervision: MB

## Funding

MB, VP, and SP are members of the ‘Sistema Nacional de Investigadores’ of the ‘Agencia Nacional de Investigación e Innovación’ (ANII) and ‘Programa de Desarrollo de las Ciencias Básicas (PEDECIBA)’ from the Universidad de la República, Uruguay.

## Acknowledgments

All authors thank Gabriela Grille for her support.

## Data Availability Statement

The original contributions presented in the study are included in the article/supplementary material, further inquiries can be directed to the corresponding author/s.

## Supplementary material

Supplementary Figure 1: Maxent modeling results.

Supplementary Figure 2: Worldwide distribution potential for R. ferrugineus obtained by Maxent modeling.

Supplementary Table 1: Supplementary_dataset_morphological_key

Supplementary Table 2: Occurrence records of R. ferrugineus Maxent.

Supplementary Table 3: Rhynchophorus spp Genbank accession numbers used to build the ML tree.

---

**Supplementary Figure 1_Figure 1:**
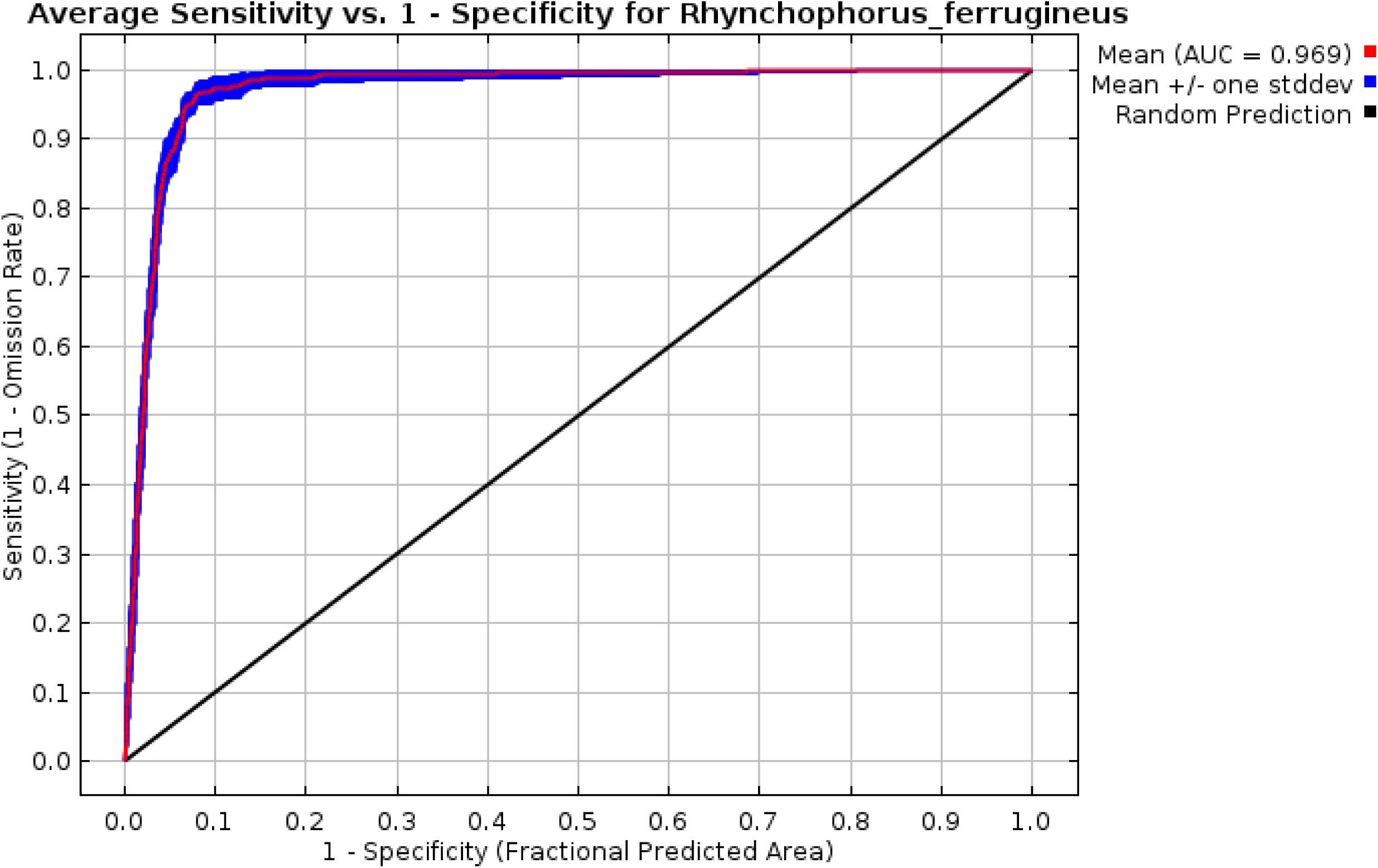
AUC result of Maxent modelling.

**Supplementary Figure 1_Figure 2:**
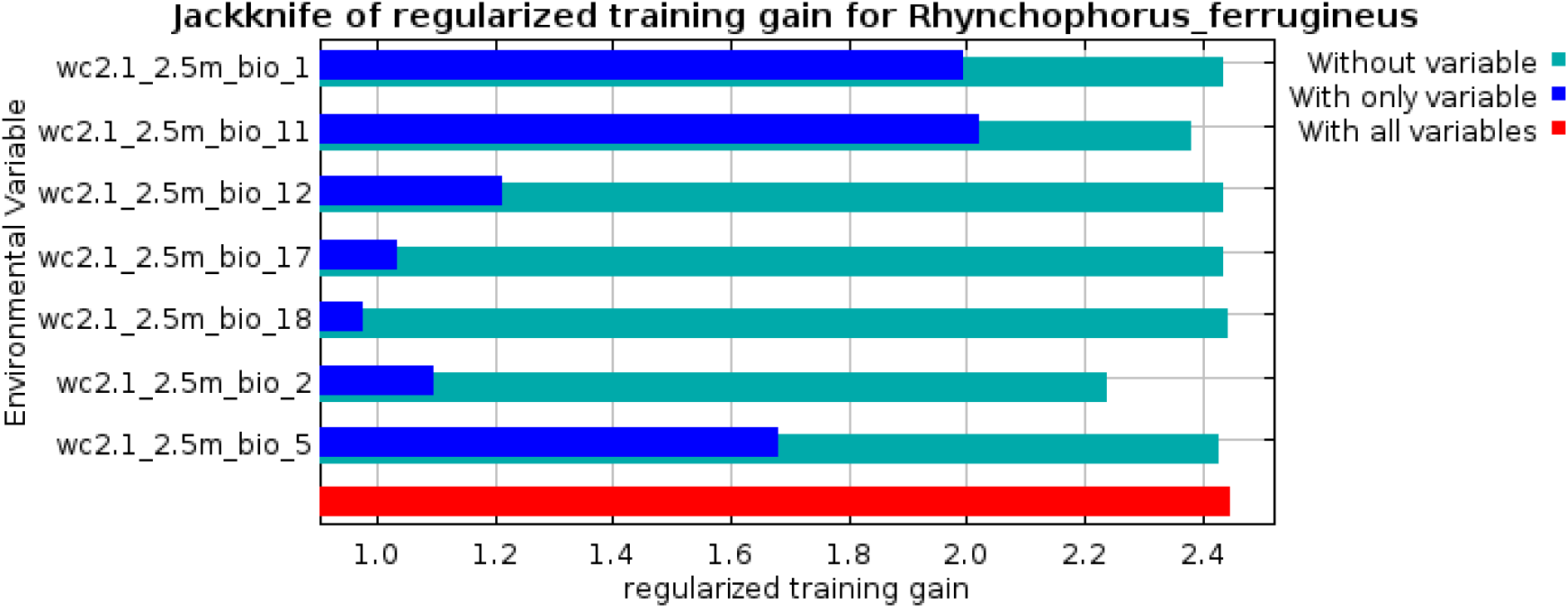
Jackknife plot of the training gain for Rhynchophorusferrugineus.

**Supplementary Figure 2:**
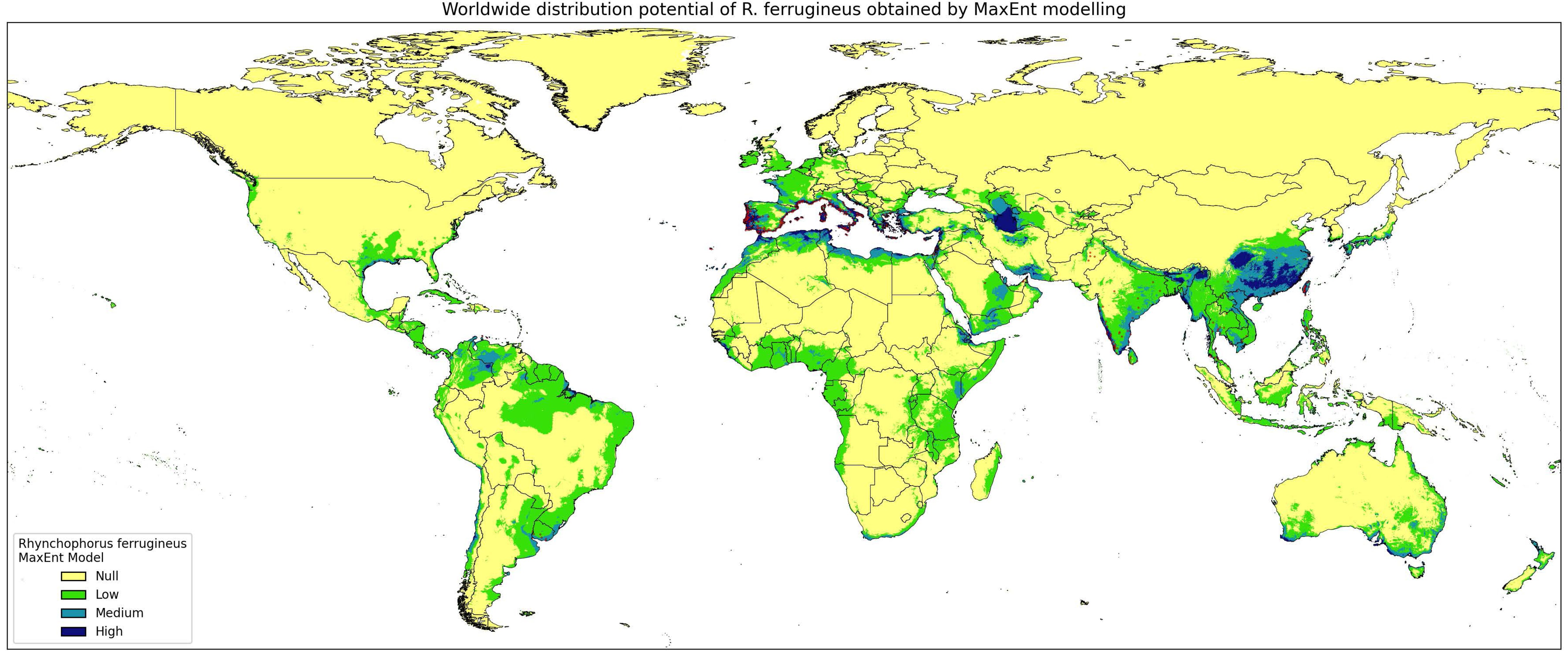
Worldwide distribution potential for *R. ferrugineus* obtained by Maxent modeling.

**Supplementary Table 1:**
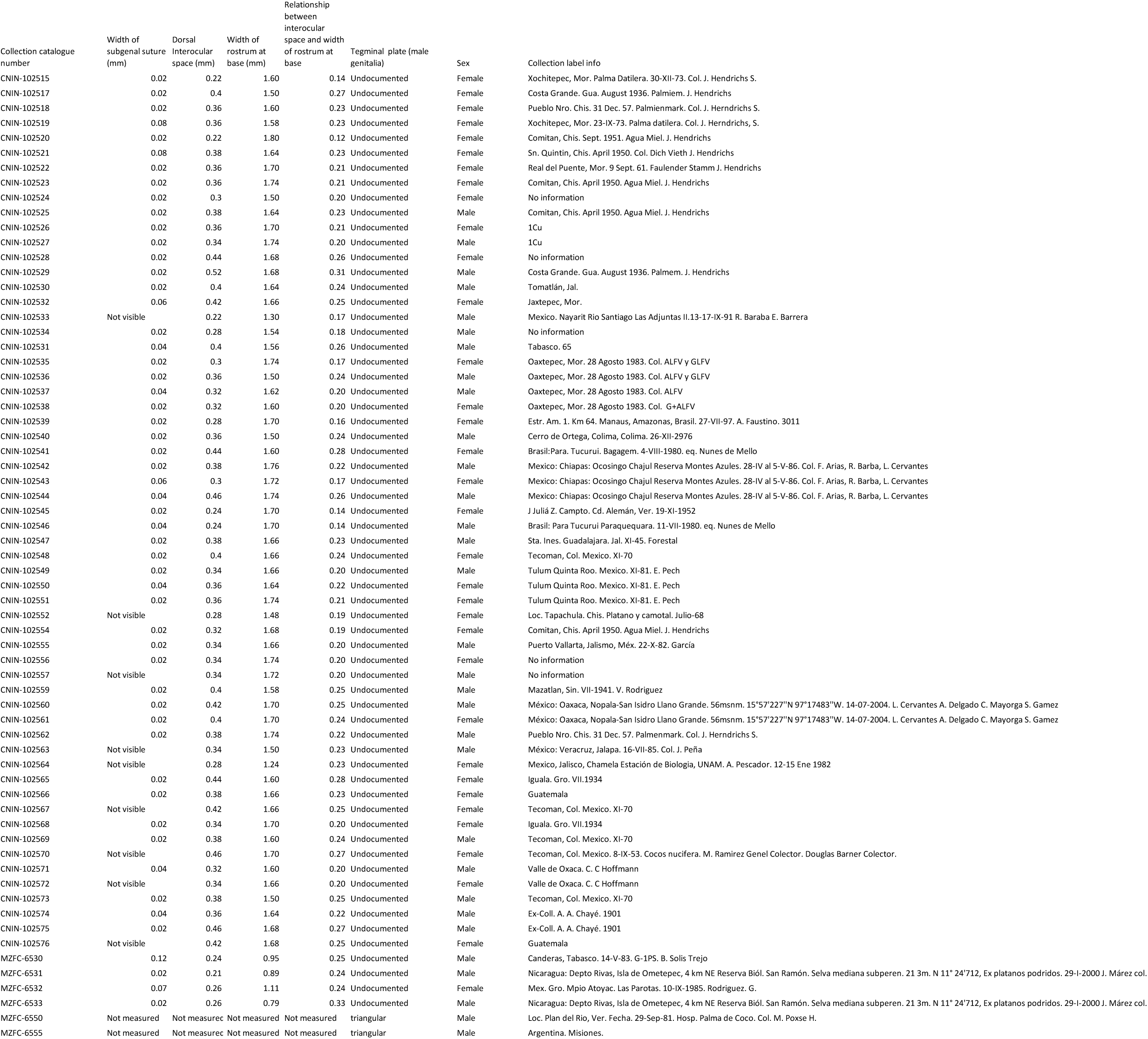
Supplementary_dataset_morphological_key.

**Supplementary Table 2:**
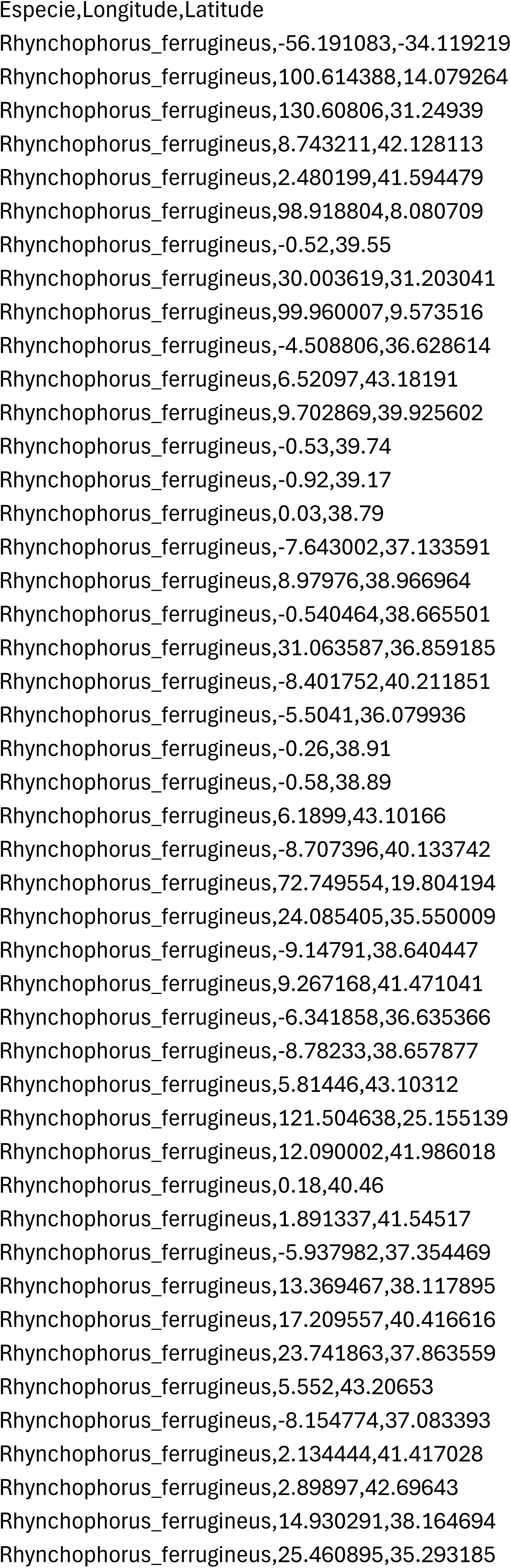
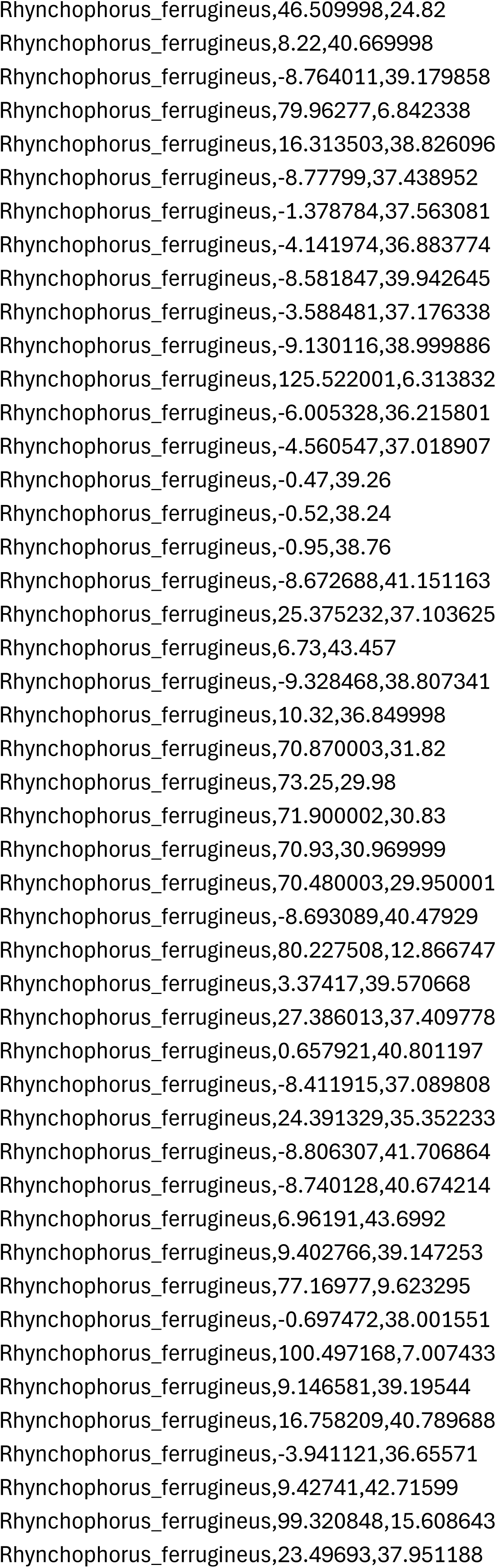
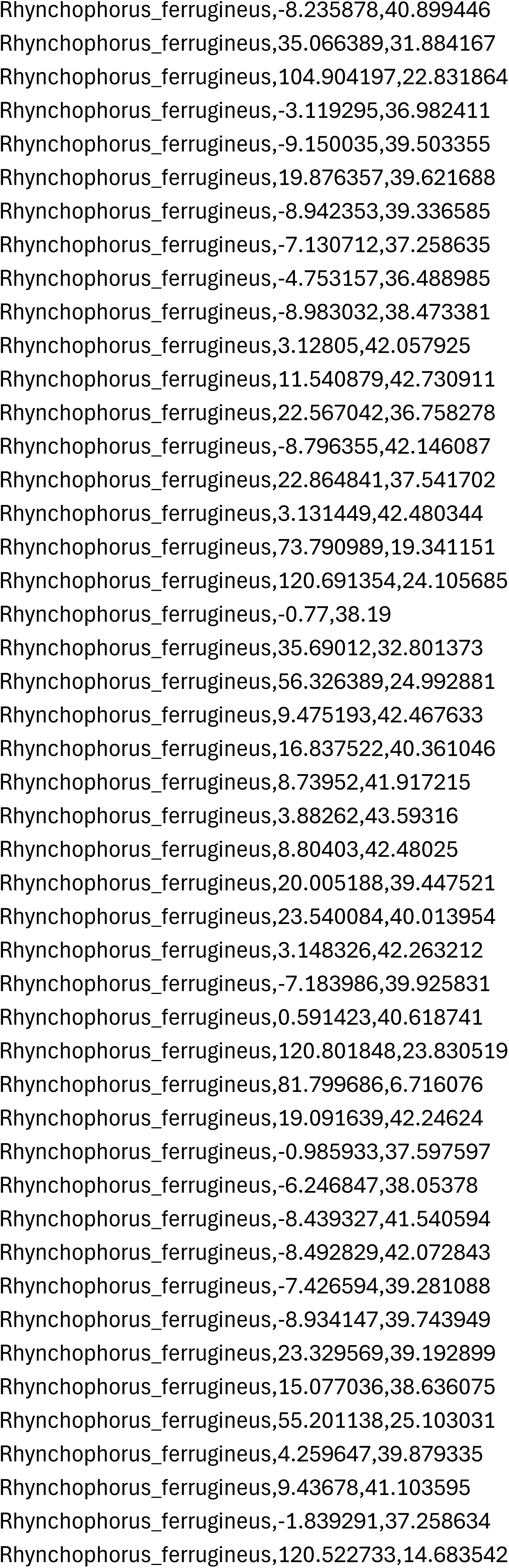
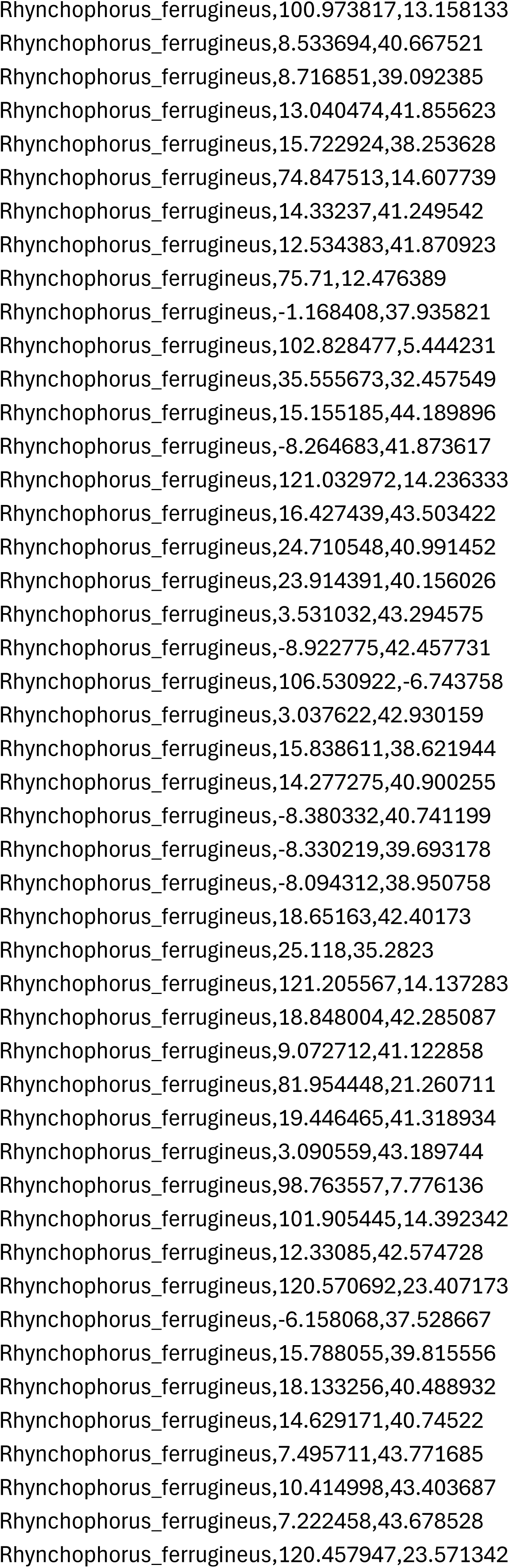
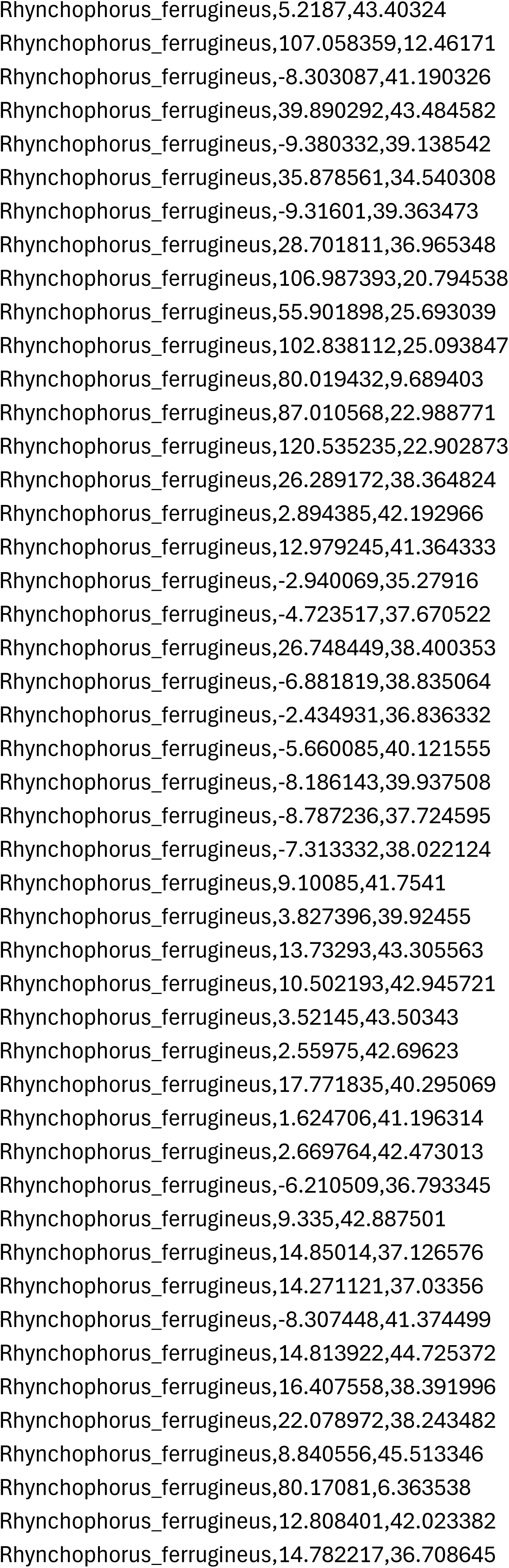
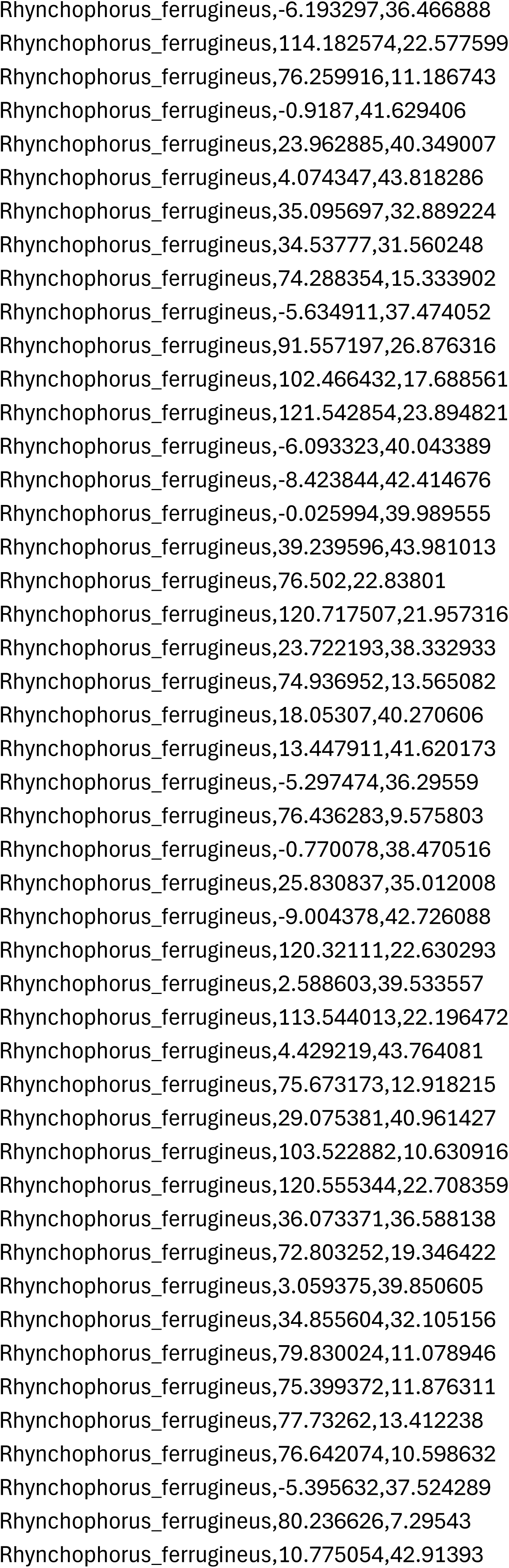
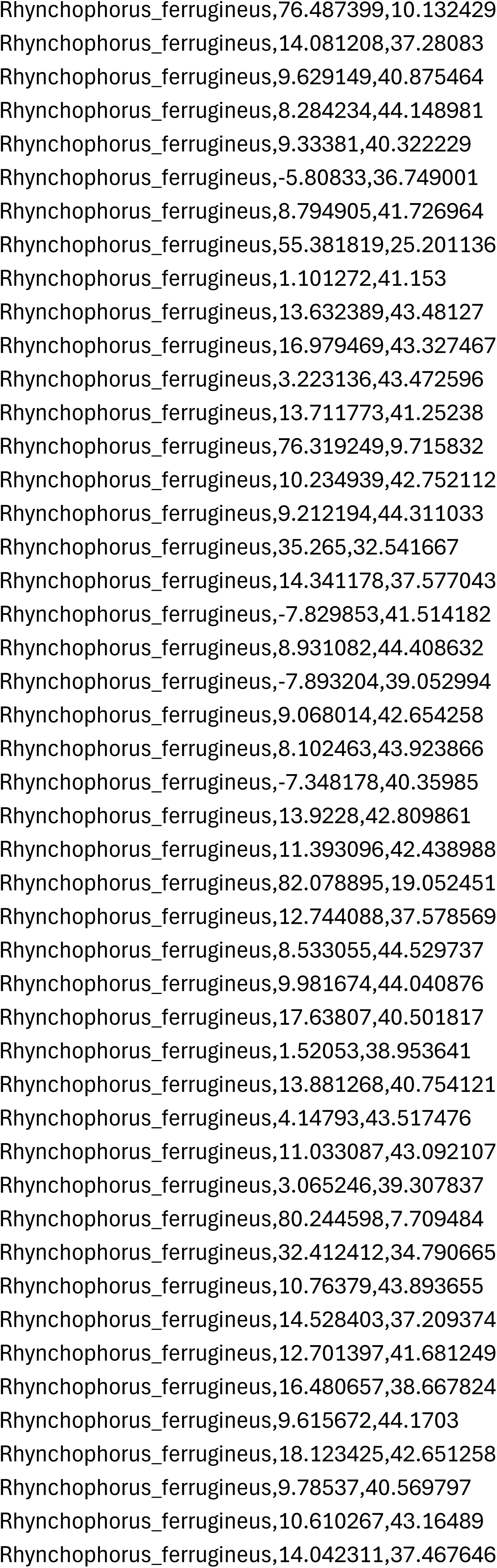
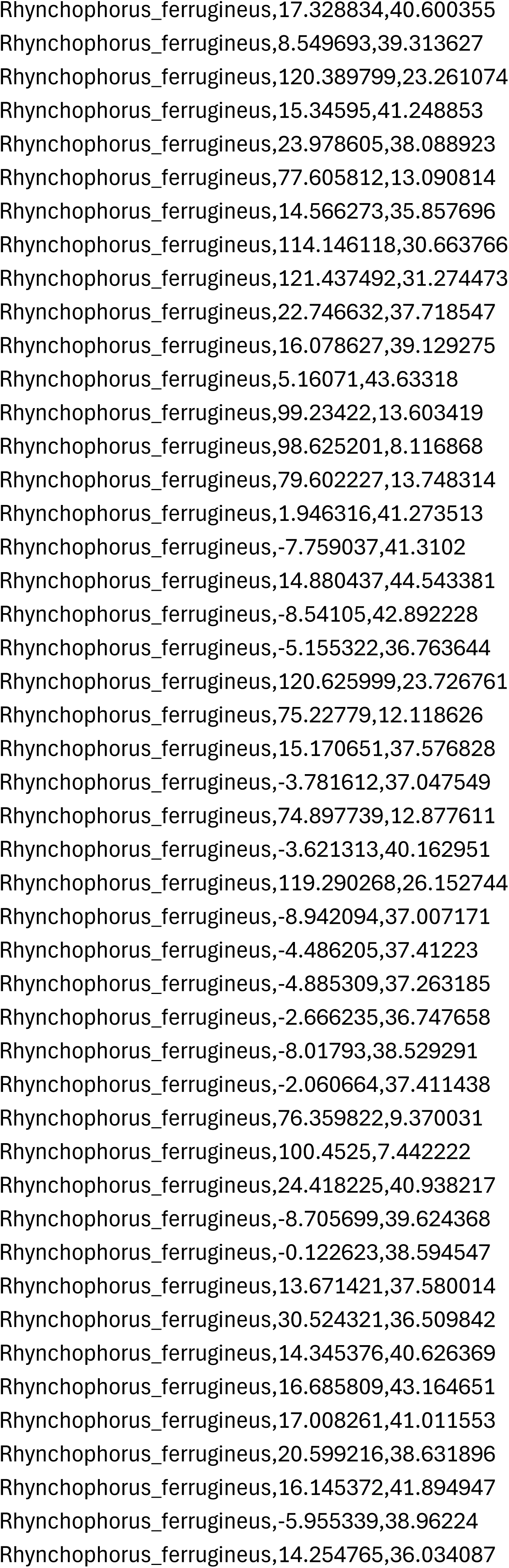
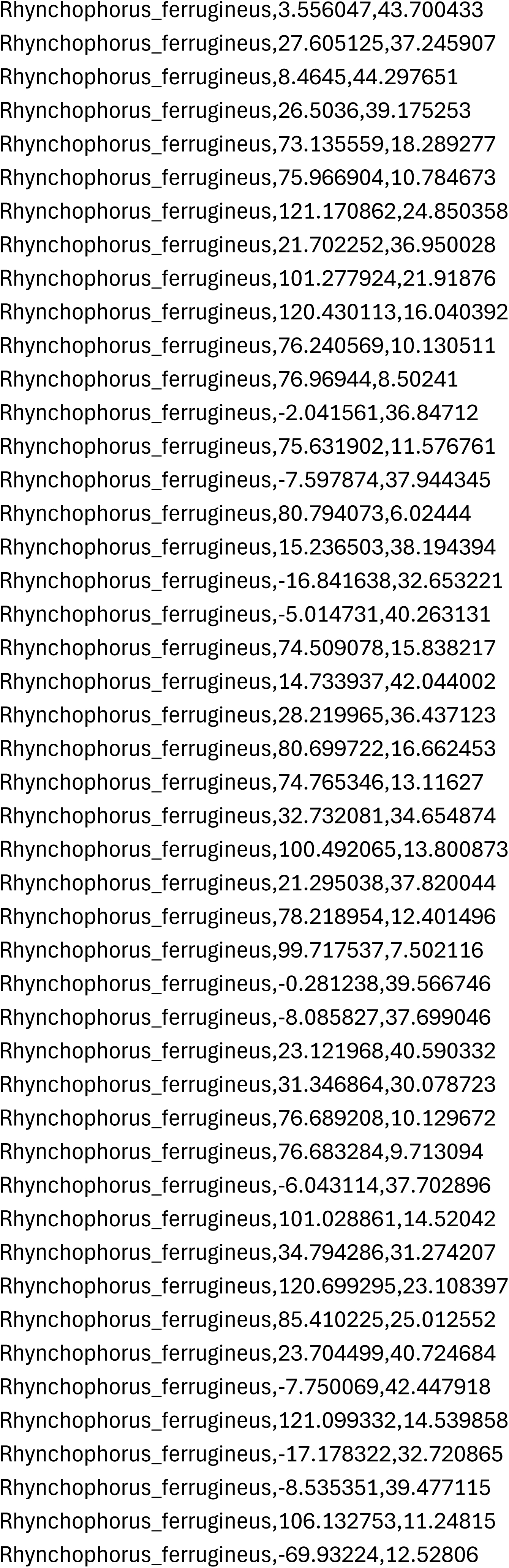
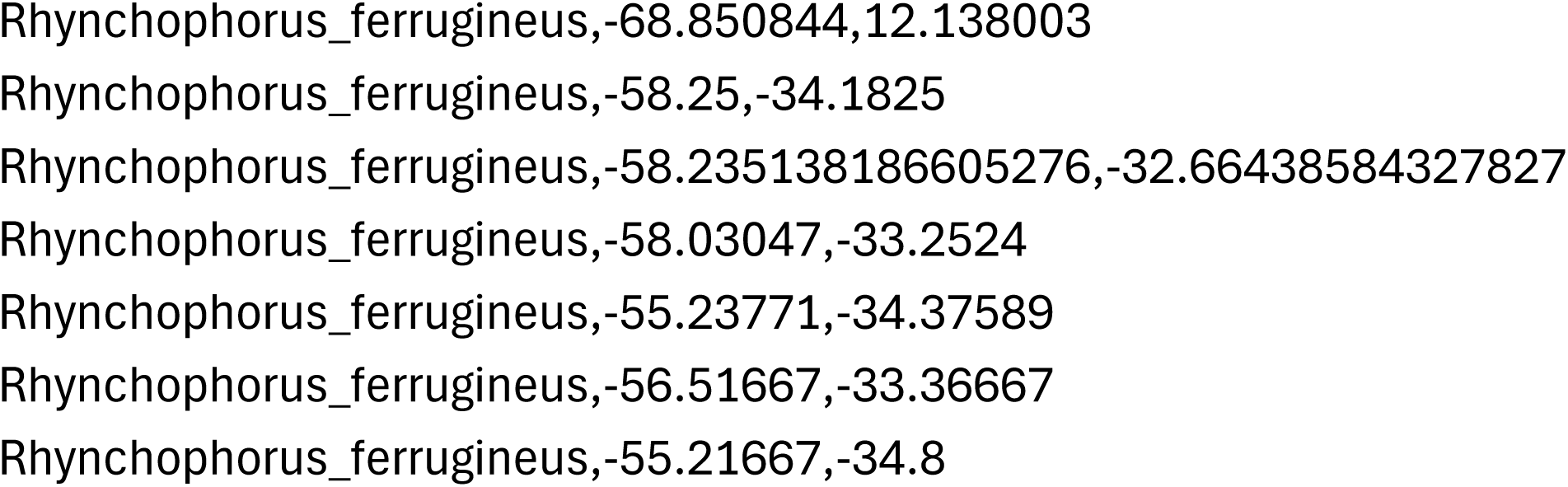
Occurrence records of *R. ferrugineus* Maxent.

**Supplementary Table 3:**
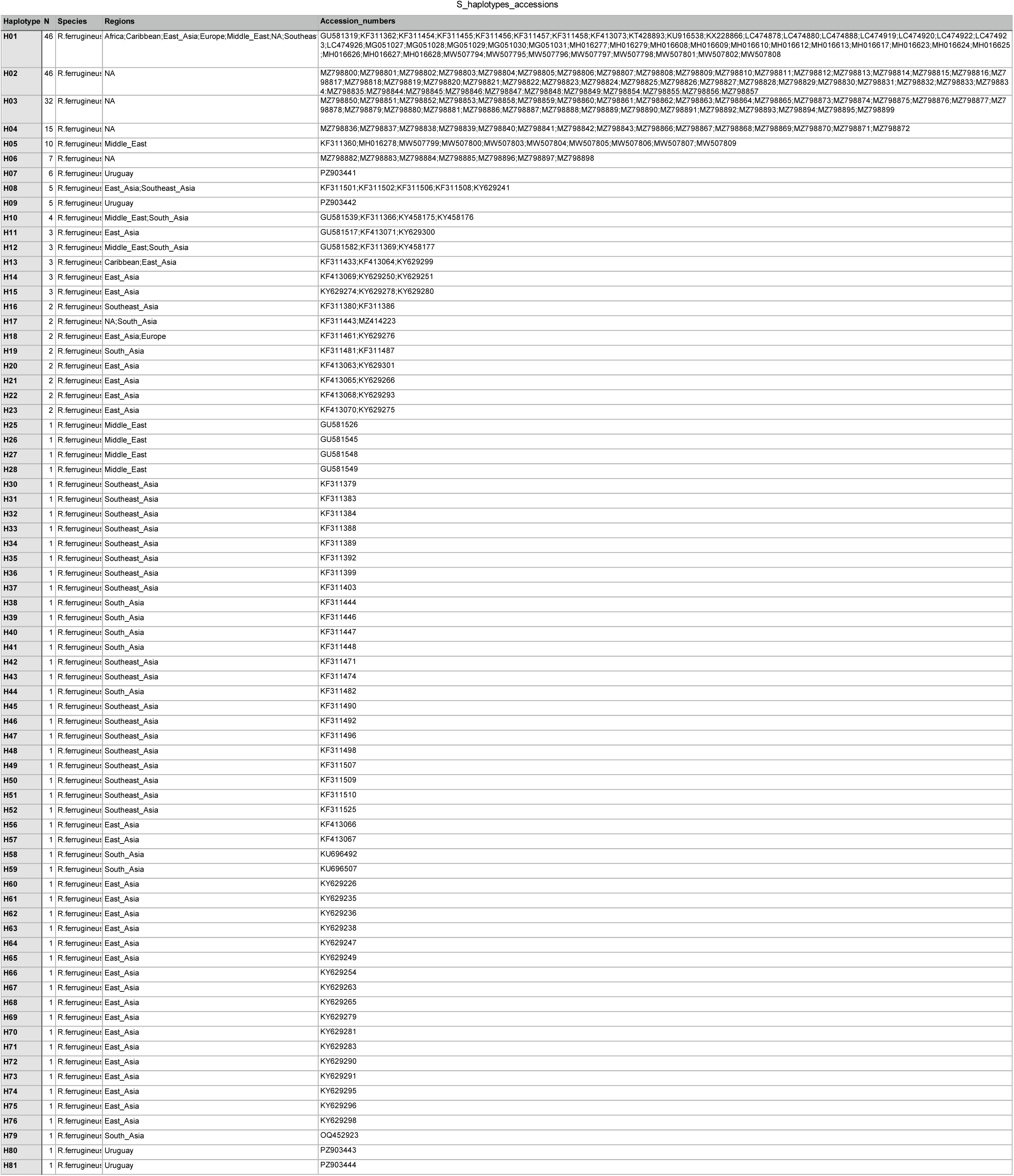
*Rhynchophorus spp* Genbank accession numbers used to build the ML tree.

**Supplementary Figure 1_Table 1:** Environmental variables used in this study and their percentage contribution (%) to variation in the data.

| Code: Variable | Percent contribution % |
| --- | --- |
| Bio 11: Mean Temperature of Coldest Quarter | 45.6 |
| Bio 01: Annual Mean Temperature | 26.0 |
| Bio 02: Mean Diurnal Range | 20.0 |
| Bio 17: Precipitation of Driest Quarter | 2.5 |
| Bio 18: Precipitation of Wettest Quarter | 2.4 |
| Bio 05: Max Temperature of Warmest Month | 2.3 |
| Bio 12: Annual Precipitation | 1.2 |

## Notes

### Competing Interest Statement

The authors have declared no competing interest.

